# Stress-Induced Transient Cell Cycle Arrest Coordinates Metabolic Resource Allocation to Balance Adaptive Tradeoffs

**DOI:** 10.1101/2020.04.08.033035

**Authors:** Alain R. Bonny, Karl Kochanowski, Maren Diether, Hana El-Samad

**Affiliations:** Department of Biochemistry and Biophysics, California Institute for Quantitative Biosciences, University of California, San Francisco, San Francisco, CA 94158, USA; Department of Pharmaceutical Chemistry, California Institute for Quantitative Biosciences, University of California, San Francisco, San Francisco, CA 94158, USA; Institute of Molecular Systems Biology, ETH Zürich, Zürich, Switzerland; Chan Zuckerberg Biohub, San Francisco, CA 94158, USA

## Abstract

The ability of a cell to mount a robust response to an environmental perturbation is paramount to its survival. While cells deploy a spectrum of specialized counter-measures to deal with stress, a near constant feature of these responses is a down regulation or arrest of the cell cycle. It has been widely assumed that this modulation of the cell cycle is instrumental in facilitating a timely response towards cellular adaptation. Here, we investigate the role of cell cycle arrest in the hyperosmotic shock response of the model organism *S. cerevisiae* by deleting the osmoshock-stabilized cell cycle inhibitor Sic1, thus enabling concurrent stress response activation and cell cycle progression. Contrary to expectation, we found that removal of stress-induced cell cycle arrest accelerated the adaptive response to osmotic shock instead of delaying it. Using a combination of time-lapse microscopy, genetic perturbations and quantitative mass spectrometry, we discovered that unabated cell cycle progression during stress enables the liquidation of internal glycogen stores, which are then shunted into the osmotic shock response to fuel a faster adaptation. Therefore, osmo-adaptation in wild type cells is delayed because cell cycle arrest diminishes the ability of the cell to tap its glycogen stores. However, acceleration of osmo-adaptation in mutant cells that do not arrest comes at the cost of acute sensitivity to a subsequent osmo-stress. This indicates that despite the ostensible advantage faster adaptation poses, there is a trade-off between the short-term benefit of faster adaptation and the vulnerability it poses to subsequent insults. We suggest that cell cycle arrest acts as a carbon flux valve to regulate the amount of material that is devoted to osmotic shock, balancing short term adaptation with long-term robustness.

## Introduction

Cells and organisms are constantly challenged in their environment with insults that vary in origin, magnitude and duration. In order to respond to these insults, cells have evolved a large battery of adaptive stress responses that allow them to survive and maintain their homeostasis. Different stress responses show a remarkable diversity in their sensing, regulation, and logic ^1–5^. However, an almost constant feature of any stress response is the involvement of cell cycle slow-down or arrest ^6–10^. It is widely assumed that this is because it is advantageous for a cell not to divide during stressful conditions in order to safeguard its own fitness and that of its future progeny ^11,12^. Furthermore, by arresting division, resources and energy can be diverted from the replication and division program to the stress response program, allowing it to proceed more efficiently ^13^. Despite the near universality of these assumptions, the precise contribution of cell cycle arrest to adaptation remains poorly understood.

Upon encountering an environmental perturbation, the model organism *Saccharomyces cerevisiae* is thought to divert its limited resources such as ribosomes to high-priority transcripts ^14,15^. However, this diversion conflicts with other ongoing and resource-intensive processes such as cell cycle progression. Concomitant cell cycle arrest can therefore relieve this resource competition, allowing the cell to adequately mitigate the effects of stress ^13^. While this model stands to reason intuitively, embedded within that conceptual framework are competing optimization problems that the cell must confront: properly addressing the stress to ensure longevity, but doing so in a time-sensitive manner for cell cycle re-entry to promote reproductive fitness. To understand the tradeoffs involved in a stress response, and interrogate the relative contribution of cell cycle arrest to cellular adaptation and intracellular process optimization, it is necessary to decouple the processes in question.

To explore this paradigm, we used the hyperosmotic glycerol (HOG) response as a convenient framework. The HOG program is a canonical stress response activated by the presence of excess osmolytes in the extracellular environment. The increase in osmotic pressure difference between the inside and outside of the cell drives water out, causing the cellular volume to decrease. At the onset of a step input of hyperosmotic shock, the central HOG mediator, Hog1, rapidly translocates from the cytoplasm to the nucleus where it interacts with a variety of targets to initiate the production and accumulation of glycerol ^16^. The accumulation of glycerol in the cytoplasm re-establishes the osmotic pressure gradient to its basal level, and once volume has been corrected, Hog1 exits the nucleus ^17^. Importantly, in addition to the initiation of glycerol synthesis, Hog1 stabilizes Sic1, the stoichiometric inhibitor of b-type cyclins 5 and 6, to transiently arrest the cell in the G1 phase of the cell cycle^11^. Volume restoration and exit of Hog1 from the nucleus also coincides with resumption of cell cycle progression ^18^. The adaptive translocation pattern of Hog1 has been the subject of many studies for its robust, reproducible and stereotyped pattern, which acts as a real-time reporter of hyperosmotic stress adaptation ^19,20^.

Using HOG as a model system, we investigated the role of transient cell cycle arrest in the adaptive response to hyperosmotic shock. Our approach was to decouple the HOG response program from the canonical cell cycle machinery by removing the stress response-cell cycle link, Sic1, such that both processes proceed simultaneously during osmotic shock. By following Hog1 translocation as a reporter of HOG adaptation, we were able to quantify deviations from Hog1’s stereotyped translocation pattern as an indication of an altered stress response. Surprisingly, we found that unabated cell cycle progression during osmoshock accelerated osmo-adaptation as measured by Hog1 translocation. Remarkably, other canonical markers of adaptation such as glycerol production and volume recovery also proceeded faster. These data indicated that cell cycle arrest impedes, rather than facilitates, adaptation to stress. To pinpoint the mechanistic roots of this phenotype, we used mass spectrometry ^13^C isotope tracing to probe the differences in metabolic flux between wild type and cell cycle arrest-disabled cells. We discovered that progression in the cell cycle during osmostress initiated catabolism of internal glycogen that is mediated by the enzyme Gph1. Breakdown of glycogen fueled faster glycerol synthesis in the mutant cells, giving them the ability to restore turgor pressure faster than the wild type. Therefore, cell cycle seems to be the guardian of a metabolic valve that remains closed when cell cycle is arrested. To investigate what vulnerabilities arise from opening of this valve, and rationalize why the wild type cells still implement cell cycle arrest despite the delay it imposes on stress adaptation, we subjected cells to repeated osmostress pulses. Under repeated pulsing, wild type cells largely return to their basal morphology while mutant cells lacking cell cycle arrest display physical traits suggestive of a compromised cell wall. This phenotype accumulates within the mutant population with each repeated stress pulse. Therefore, adaptation to an osmotic stress proceeds faster when there is no cell cycle arrest, but this phenotype leaves the cells particularly susceptible to subsequent osmoshocks. Our findings reveal an example where connection between three important cellular networks - a stress pathway, cell cycle regulation, and metabolic control - collaborate in order to strike a balance between mounting a rapid adaptive response to acute threats and prioritizing robustness in the face of future insults.

## Results

### Removal of Hog1-mediated cell cycle arrest accelerates adaptation to hyperosmotic shock

To assess the role of cell cycle arrest in adaptation to osmotic shock, we removed the ability of Hog1 to initiate cell cycle arrest by generating a Sic1 knockout strain (Figure 1A). In this strain, we tagged Hog1 with mVenus at its endogenous locus to allow visualization of its nuclear translocation by microscopy. We also incorporated the same Hog1-mVenus construct in a wild type (WT) strain to allow for comparison of its Hog1 dynamics with those of the *sic1Δ* mutant. For precise temporal control in applying a step input of osmotic shock, we used a commercially available microfluidic platform that allowed us to quickly induce osmoshock and monitor Hog1 dynamics in the *sic1Δ* and WT strains by time lapse microscopy. We chose the osmolyte sorbitol as the input to induce hyper-osmotic stress because it is an inert sugar in the presence of glucose ^21^.

**Figure 1:**
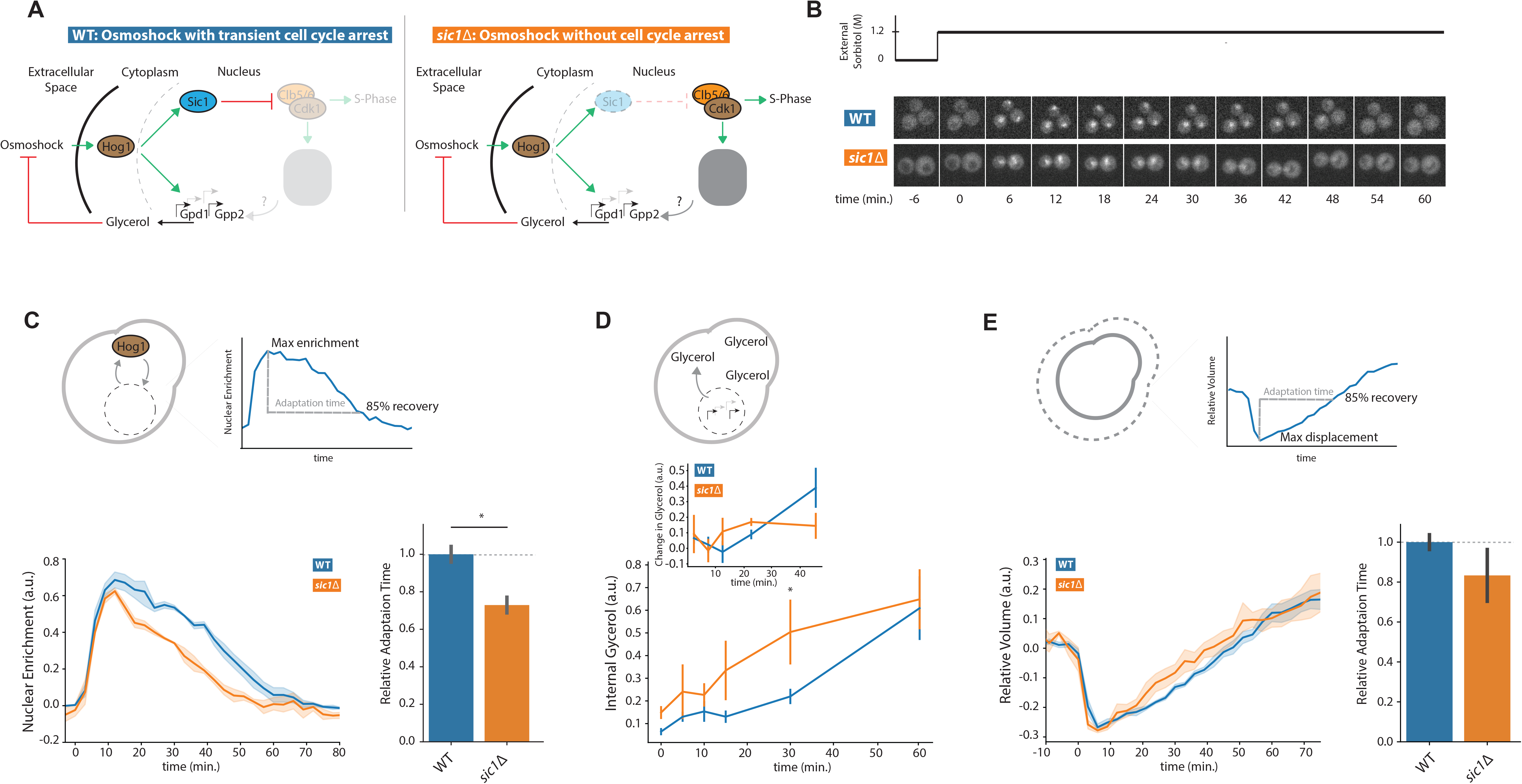
Removing cell cycle arrest by deletion of Sic1 accelerates the HOG adaptation program during osmotic shock. A) A simplified schematic of the HOG pathway showing its coupling to cell cycle arrest and glycerol production. In a cell cycle mutant strain *sic1Δ*, we asked whether the removal of stress-induced cell cycle arrest affects the adaptive response. B) Representative time lapse images of endogenously-tagged Hog1-mVenus following step input of 1.2 M sorbitol osmotic shock to WT (top) and *sic1Δ* (bottom) cells. Quantification of all cells presented in Panel C. C) Top: Cartoon depicting the translocation dynamics and quantification of adaptation time. Bottom: Quantification of the WT (blue) and *sic1Δ* (orange) Hog1-mVenus nuclear enrichment in the experiment described in Panel B. Shaded regions represent the standard error of the mean (SEM) of n=3 biological replicates. Right: Quantification of Hog1 adaptation for WT (blue) and *sic1Δ* (orange). Values are normalized to the average of WT. Error bars represent the SEM of n=3 biological replicates. *P-value<0.05; two-sided Student’s *t*-test. D) Top: Cartoon schematic depicting the intracellular accumulation of glycerol. Bottom: Quantification of internal glycerol as a function of time for WT (blue) and *sic1Δ* (orange) to a step input of 1.2 M sorbitol osmotic shock. Measurements are taken using a colorimetric assay. Error bars represent the standard deviation for n=3 biological replicates. Inset: the change in glycerol, calculated as the difference between two time points for data in Panel D, is plotted as a function of time. *P-value<0.05; two-sided Student’s *t*-test. E) Top: Cartoon schematic depicting volume recovery and quantification of its adaptation time. Bottom: Change in volume of the WT (blue) and *sic1Δ* (orange) strains in response to a step input of 1.2 M sorbitol osmotic shock. Shaded regions represent the SEM of n=4 biological replicates. Right: Quantification of volume adaptation time of WT (blue) and *sic1Δ* (orange) volume. Values are normalized to the average of WT. Error bars represent the SEM of n=4 biological replicates.

Using this experimental setup, we subjected WT and *sic1Δ* mutant cells to a step input of 1.2 M sorbitol osmoshock and monitored Hog1 nuclear translocation dynamics. Following the sorbitol input, Hog1 rapidly translocated to the nucleus in both strains with similar nuclear influx dynamics (Figure 1B). The degree of maximum nuclear enrichment was virtually indistinguishable in both strains, suggesting that the mutant maintains the ability to sense and respond to acute osmotic shock. In the WT strain, Hog1 exited the nucleus and its cytoplasmic levels adapted to the pre-stimulus values within 45 minutes on average, consistent with previous reports ^20^. Surprisingly, however, the return of Hog1 to the cytoplasm was much faster in the *sic1Δ* cells, occurring on average within 33 minutes (Figure 1C). This constitutes a significant 30% speed-up compared to WT (Figure 1C, right). This acceleration of Hog1 adaptation in non-arresting cells was not a sorbitol-specific effect, since experiments carried out with 0.6 M NaCl also showed the same phenotype (Supplemental Figure 1A, B). The Sic1 protein is known to be a G1-specific regulator^22^. However, since we observe this phenotype averaged over the entire asynchronous population, we are likely underestimating the effect of its deletion.

The faster Hog1 response in *sic1Δ* cells can be the result of a breakdown of coordination between the regulatory osmotic response and turgor pressure of the cell. If this were the case, then Hog1 would recover its cytoplasmic localization without full recovery in other physiological parameters such as cellular volume and glycerol accumulation necessary for this recovery. On the other hand, if the fast Hog1 adaptation were the result of an acceleration within an intact recovery program, then the profile of volume recovery and glycerol accumulation should also be accelerated. We therefore investigated these critical phenotypes to see if the integrity of the adaptive program is maintained despite accelerated Hog1 dynamics in the *sic1Δ* strain.

First, we measured internal glycerol content using a colorimetric assay following a 1.2M sorbitol input over 60 minutes in both the WT and *sic1*Δ strains (Figure 1D). Using this assay, we determined that both WT and mutant cells upregulated their glycerol production by approximately 5-fold at the end of the 60 minutes. However, while the WT cells did not begin to dramatically upregulate glycerol production until 30 minutes after the onset of stress, *sic1*Δ began increasing glycerol synthesis only 15 minutes after stress (Figure 1D, inset). In agreement with this finding, quantification of cellular surface area as a surrogate for volume also showed a faster recovery for the mutant relative to the WT (Figure 1E). While this volume recovery phenotype was reproducible, its extent was slightly lower than the Hog1 and glycerol phenotype, showing ~15% difference between WT and mutant (Figure 1E, right). This could be due to the difficulty in cell tracking and quantification of surface area, or to a strict upper limit on the expansion properties of the cell wall ^23^. Taken together, the three phenotypes strongly support the hypothesis that the mutant has the same coordinated osmotic stress response as the WT type, but with faster dynamics that ensue from the inability of these cells to arrest their cell cycle during osmotic shock.

### Glycerol production is accelerated by using internal sources in mutant that lacks cell cycle arrest

Because Hog1 translocation, glycerol production and volume all correlate with faster recovery dynamics in the mutant strain, we next sought to investigate if another cellular process was fueling the accelerated production of glycerol, the effector molecule for osmotic shock recovery. We hypothesized that in the *sic1*Δ strain, a surplus of carbon material from central glycolysis could be shunted into glycerol production, thus resulting in heightened glycerol synthesis.

One possible scenario for this to happen is one in which the deletion of Sic1 augments the ability of cells to import extracellular glucose, resulting in a greater amount of carbon material entering glycolysis to be directed towards glycerol production. To test this scenario we performed a mass spectrometry ^13^C isotope tracing experiment to compare the external glucose incorporation rate between the mutant and WT strains. In this experiment, cells were grown in ^12^C glucose media. At time zero, an aliquot of cells was transferred onto filter paper over a vacuum manifold and continuously perfused with fully-labeled ^13^C glucose media with and without 1.2 M sorbitol for various durations before quenching the sample (Figure 2A, Supplemental Figure 2A). A pilot experiment (not shown) suggested that turnover rates of glycolytic intermediates occur on the order of seconds, with intermediates reaching steady-state after 1 minute. Therefore, we chose to quench samples at 10, 20, 30, 45 and 60 seconds in order to capture the rate at which the internally ^12^C-enriched glycolysis intermediates are degraded and newly synthetized metabolites incorporate the perfused ^13^C. Consistent with a fast turn-over of these metabolites, we observed a rapid decay of ^12^C enrichment among the glycolysis intermediates within seconds. In the examples of glucose-6-phosphate (G6P) and fructose bisphosphate (FBP), the *sic1Δ* strain incorporated ^13^C with a slower rate than the WT in the presence and absence of osmotic shock (Figure 2B, Supplemental Figure 2B). Using the decay rate of ^12^C enrichment as a surrogate for extracellular glucose incorporation, we derived ^13^C incorporation rates of 0.117 s^−1^ and 0.04 s^−1^ for WT and *sic1Δ*, respectively, during osmotic shock for G6P. Similarly, FBP incorporated ^13^C at rates of 0.12 s^−1^ and 0.064 s^−1^ for WT and *sic1Δ*, respectively. The approximately 2-to-3-fold slower ^13^C incorporation rate in the *sic1Δ* mutant was consistent for all glycolysis intermediates we targeted, from upper glycolysis in G6P to phosphoenolpyruvate (PEP) in lower glycolysis (Supplemental Figure 2B,C). Therefore, it is clear that the faster recovery of the *sic1Δ* mutant is unlikely to be ascribed to import of external glucose, since the mutant is slower at ^13^C incorporation.

**Figure 2:**
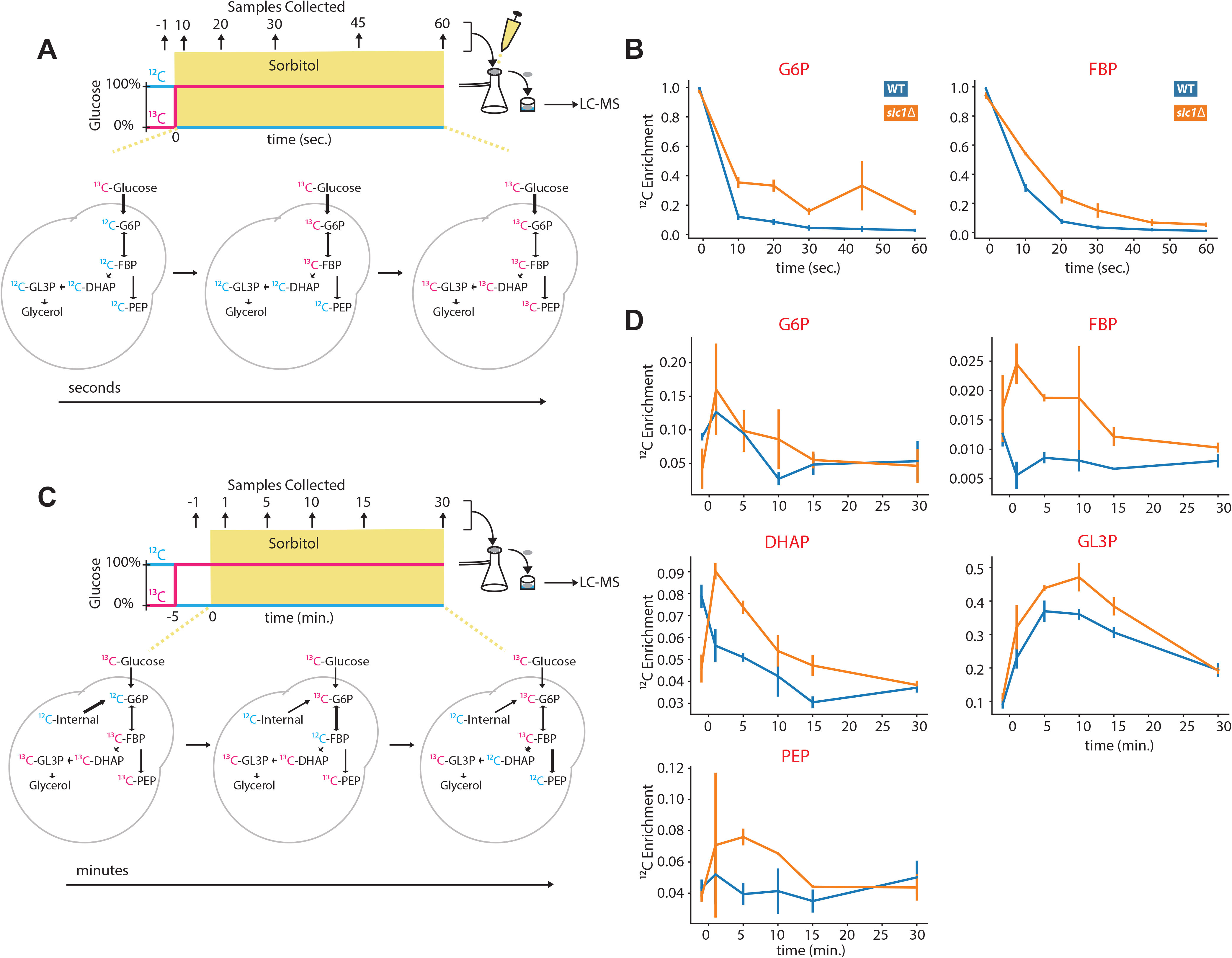
An internal carbon store is shunted towards excess glycerol production during osmotic shock in the *sic1Δ* mutant. A) Cartoon schematic of the experiment to measure extracellular glucose incorporation rates. Cells were inoculated overnight, diluted and outgrown in ^12^C glucose. At time zero a 1 mL sample of cells was transferred to filter paper above a vacuum manifold and continuously perfused with fully-labeled ^13^C glucose media. A 1.2 M sorbitol input was also administered at time 0. Samples were taken at 10 s, 20 s, 30 s, 45 s, and 60 s and transferred to quenching solution. B) ^12^C enrichment over time of central glycolysis metabolite glucose-6-phosphate (G6P) (Left) and fructose bisphosphate (FBP) (Right). Traces shown are WT (blue) and *sic1Δ* (orange) strains for experiment described in Panel A. Error bars represent the standard deviation of n=2 technical replicates. C) Cartoon schematic of experiment to test internal carbon enrichment of targeted metabolites. Cells were inoculated overnight, diluted and outgrown in ^12^C glucose. Five minutes prior to time zero, cells were resuspended in fully-labeled ^13^C glucose. At time zero the culture was diluted 1:1 with 2.4 M sorbitol in fully-labeled in ^13^C glucose. At the indicated time points, 1 mL of culture was placed on filter paper above a vacuum manifold for the media to wash through, transferred to quenching solution and then measured. D) ^12^C enrichment over time for a panel of select metabolites in glycolysis and glycerol production. Traces shown are the WT (blue) and *sic1Δ* (orange) strains for experiment described in Panel C. Error bars represent the standard deviation of n=2 technical replicates.

Slower glucose incorporation often has the implication of a reduced doubling rate, which has been suggested to confer an advantage during stress adaptation ^24,25^. We measured the growth rate of the *sic1Δ* mutant and observed that it has a doubling time of 3.4 hours compared to 2.1 hours of the WT in defined media (Supplemental Figure 3A). To assess whether slower growth could be correlated to the accelerated osmoshock adaptation, we compared the growth rate of all strains used in this study (see subsequent sections for different strains, including those with different deletions in metabolic genes) against their respective adaptation time to an osmotic shock (Supplemental Figure 3B). We found a weak correlation (R =0.34) between the growth rate of the strains tested and their adaptation time to osmotic stress, suggesting that while growth may contribute to the accelerated phenotype, it is unlikely to be the only factor.

The difference in external glucose incorporation rates between WT and *sic1Δ* suggest that the mutant is not importing more extracellular carbon material into glycolysis. To further investigate the source of its accelerated glycerol production, we reasoned that if the excess carbon material was not from extracellular sources, then it was conceivable that the glycerol synthesis in the *sic1Δ* strain could be assisted by diverted intracellular carbon stores. To investigate this possibility, we again turned to ^13^C mass spectrometry isotope tracing to test whether, and to what extent, intracellular carbon was used for glycerol production in both strains. In order to selectively enrich internal stores with a unique carbon isotope, we grew the cells on ^12^C glucose, and 5 minutes before time zero, resuspended the cells in fully-labeled ^13^C glucose. We then continued this treatment with and without osmoshock (1.2M sorbitol) for the remainder of the experiment, collecting samples until 30 minutes after osmoshock (Figure 2C). Based on the rapid turn-over of glycolytic intermediates (Supplemental Figure 2B, C), we reasoned that within 5 minutes all glycolysis intermediates will be ^13^C enriched, but macromolecules such as storage carbohydrates would remain enriched in ^12^C due to their slower turn-over ^26,27^. Given that the cell is only provided ^13^C carbon at the onset of osmotic shock, and glycolysis intermediates likely enriched ^13^C in the 5 minutes before stress, any detection of ^12^C after HOG activation would implicate the liquidation of an internal macromolecule.

By monitoring the panel of targeted metabolites in glycolysis and glycerol precursors, we detected low ^12^C enrichment along central glycolysis, suggesting that both strains liquidated internal stores during osmoshock. The dynamic pattern of ^12^C enrichment of metabolites for the WT cells showed a pattern of low and unchanged level (FBP and PEP) or a level that declined as a function of time (G6P and DHAP) (Figure 2D, Supplemental Figure 4B). The temporal enrichment dynamics of the *sic1Δ* mutant, however, were markedly different showing a transient peak of ^12^C enrichment (indicating incorporation of internal carbon stores) followed by a decrease for all metabolites (Figure 2D, Supplemental Figure 4B). In the first step of glycolysis, the metabolite G6P started with a low ^12^C enrichment, which increased and peaked between 1 and 5 minutes and subsequently decreased. The same pattern was present for glycolysis intermediates FBP and PEP. More importantly, this same transient ^12^C signature was present in the glycerol production branch represented by dihydroxyacetone phosphate (DHAP). This transient ^12^C enrichment pattern in the *sic1Δ* mutant suggests that ^13^C in glycolysis intermediates is temporarily replaced by ^12^C from internal stores in a flux that traverses the metabolic route to glycerol production. However, after this wave, external ^13^C is incorporated again in metabolites, underlying the subsequent decline in ^12^C enrichment.

**Figure 4:**
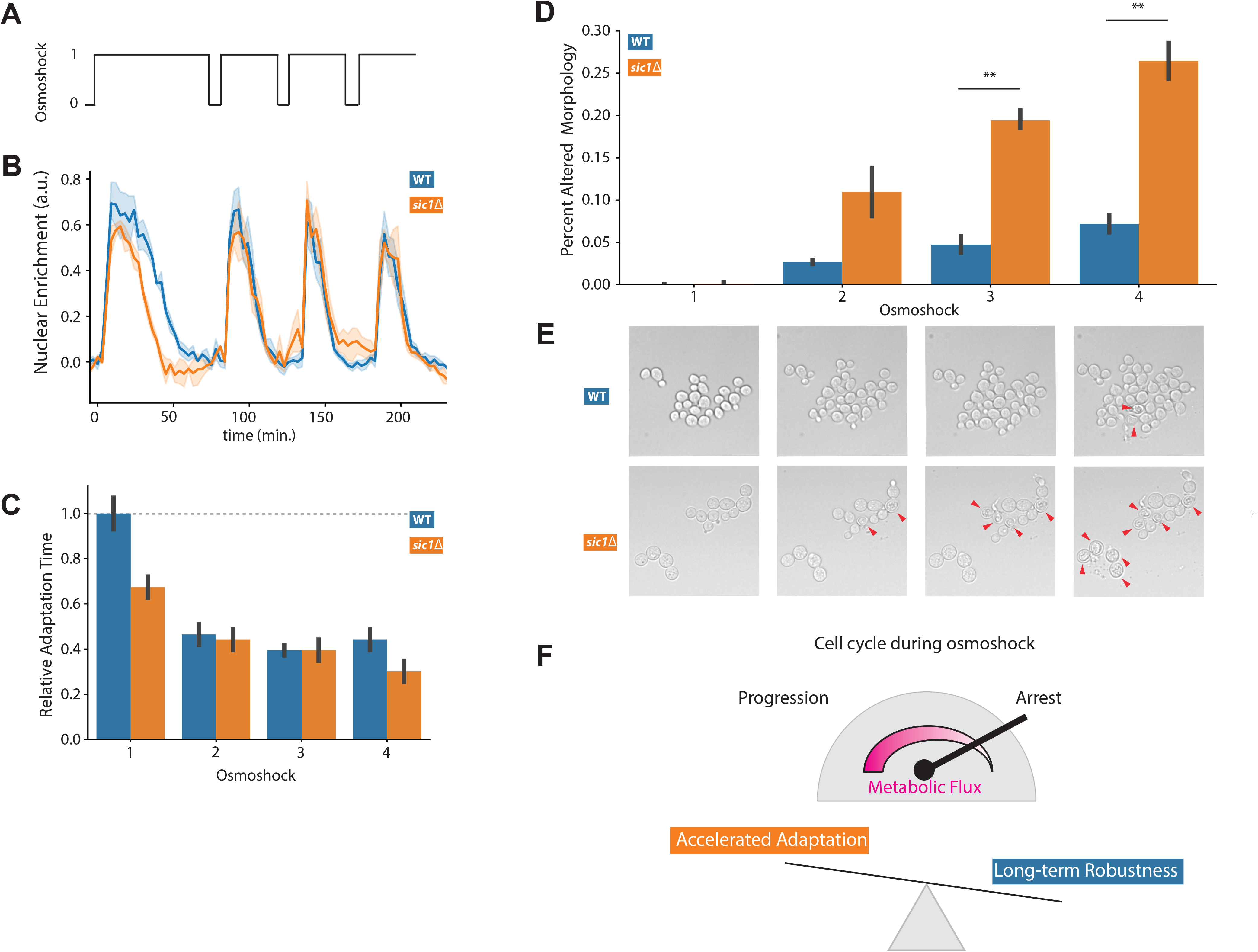
Cell cycle arrest mediates tradeoffs between fast recovery and resilience to multiple instances of osmotic shock. A) Top: experiment schematic representing a series of 1.2 M sorbitol osmotic shock step inputs. The first input lasts for 90 minutes and subsequent inputs last 45 minutes, and are separated by 5 minutes. B) Time traces of Hog1 nuclear enrichment of WT (blue), *sic1Δ* (orange). Shaded regions represent the SEM of n=3 biological replicates. C) Quantification of adaptation time of Hog1 nuclear enrichment for WT and *sic1Δ* strains for data presented in Panel B. Values are normalized to the average first response for the WT strain. Error bars represent the SEM of n=3 biological replicates. D) The percent of cells with altered morphological phenotypes at the end of each step input of 1.2 M sorbitol. Error bars represent the SEM of n=3 biological replicates. **P<0.005; two-sided Student’s *t*-test. E) Representative brightfield images corresponding to the timepoints quantified in Panel D. Images depict the compromised cell wall morphology indicated by red arrows in the WT (top) and *sic1Δ* (bottom) strains. F) Conceptual model of the role of cell cycle arrest in hyperosmotic shock response.

It is worth noting that in the absence of stress, the WT had a higher basal ^12^C enrichment across the metabolites targeted, likely due to a higher basal turnover rate of macromolecules, and consistent with its faster growth rate compared to the *sic1Δ* strain ^26^. Only during the onset of stress did the *sic1Δ* mutant have a brief, higher ^12^C enrichment with the aforementioned peak throughout (Figure 2C, Supplemental Figure 4B). Collectively, these data strongly suggest that, unlike the WT, *sic1Δ* briefly shunts internal carbon stores as extra flux into glycerol production at the beginning of osmotic shock.

Interestingly, the immediate precursor to glycerol, glycerol-3-phosphate (GL3P), had an order of magnitude greater proportion of ^12^C compared to the other metabolites tested (Figure 2D). The increase in ^12^C could be attributed to back-flux from existing ^12^C-enriched glycerol by way of a futile cycle to degrade excess ATP during severe stress ^28^. Despite the discrepancy between GL3P and DHAP, the *sic1Δ* strain was still enriched with a greater amount of ^12^C than the WT, likely reflecting the convergence of the aforementioned futile cycle and internal carbon liquidation.

### Internal glycogen is liquidated using the Gph1 enzyme and shunted into glycerol production in mutant that lacks cell cycle arrest

Previous studies have established links between cell cycle progression and central metabolism as integral to cellular physiology ^29–31^. These links are mediated mechanistically through biochemical interactions where CDK1 (Cdc28) activates storage catabolism enzymes for trehalose and glycogen (Nth1 and Gph1, respectively) ^32,33^. Given these data, we hypothesized that during osmotic shock, the *sic1Δ* mutant could activate storage catabolism enzymes through unabated cell cycle progression, resulting in a burst of glycolytic flux that was then shunted into excess glycerol production (Figure 3A). Further, we predicted that by coupling *sic1Δ* with either a Nth1 or Gph1 knockout, we could rescue the mutant to the WT phenotype, for example as measured by Hog1 localization dynamics.

**Figure 3:**
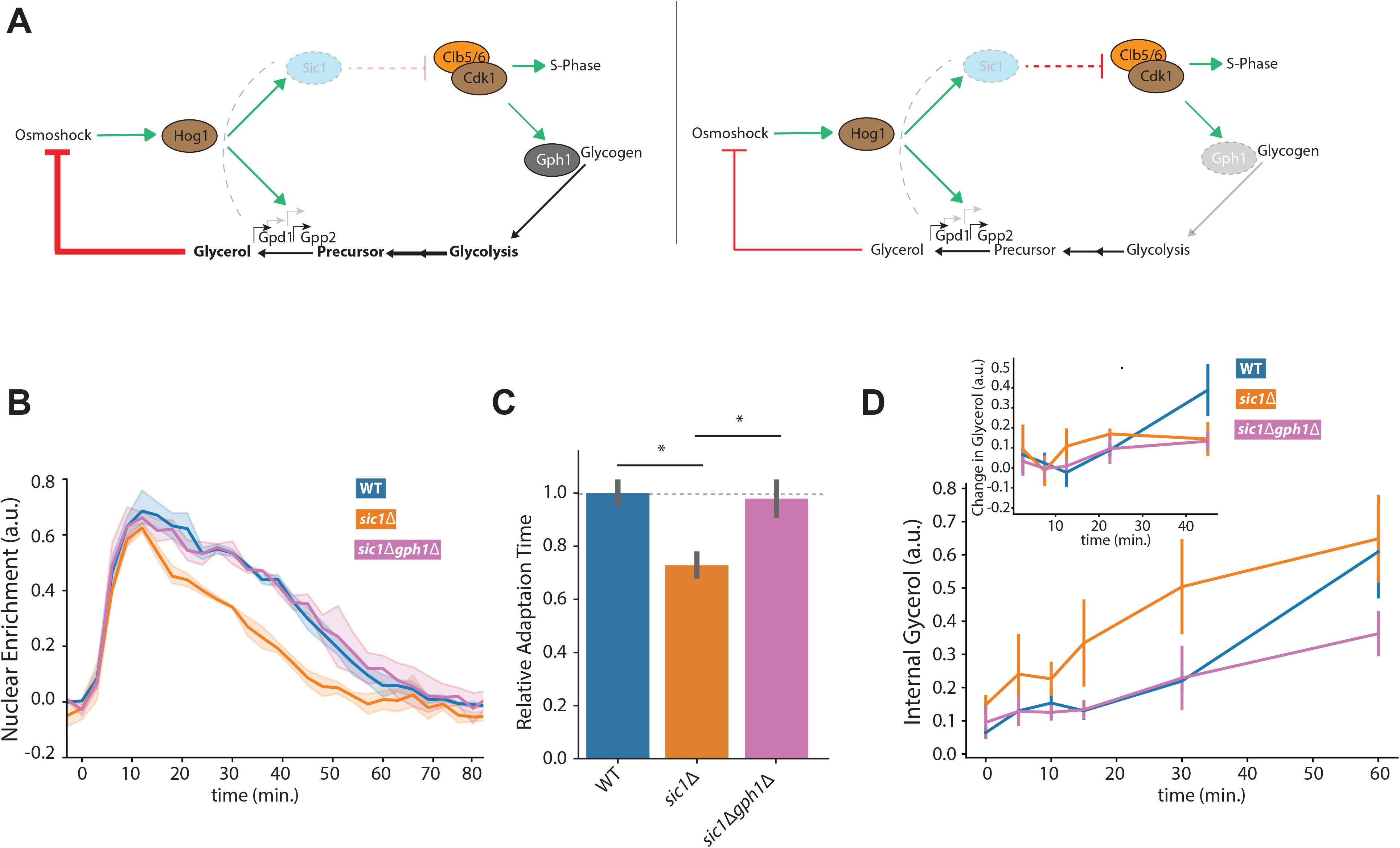
Glycogen catabolism enzyme Gph1 mediates expedited glycerol synthesis to fuel acceleration phenotype in *sic1Δ* mutant. A) Schematic depicting the hypothesis that liquidation of glycogen by activation of Gph1 accelerates the Hog1-mediated glycerol production. B) Left: Traces of Hog1 nuclear enrichment over time following 1.2 M sorbitol osmotic shock in the WT (blue), *sic1Δ* (orange), *sic1Δgph1Δ* (purple) cells. Shaded regions represent the SEM of n=3 biological replicates. C) Quantification of adaptation time of Hog1 nuclear enrichment computed as in Figure 1B. Values are normalized to the average WT. Error bars represent the SEM of n=3 biological replicates. *P<0.05; two-sided Student’s *t*-test. D) Measurement of internal glycerol over time for the strains shown in Panel B in response to a step input of 1.2 M sorbitol osmotic shock. Measurements are taken using a colorimetric assay. Inset: the change in glycerol, calculated as the difference between two time points for data in Panel D, is plotted as a function of time. Error bars represent the standard deviation for n=3 biological replicates.

Cellular trehalose levels have been widely established as mediators of stress recovery ^34^. However, surprisingly, in a *sic1Δnth1Δ* mutant, Hog1 nuclear levels still adapted significantly faster than in WT following osmotic stress (Supplemental Figure 5A, B), suggesting that trehalose is not the main liquidated internal carbon source. However, when we coupled the *sic1Δ* deletion with a knockout of Gph1 (*sic1Δgph1Δ* mutant), Hog1 adapted nearly 30% slower, closely resembling WT recovery time (Figure 3B, C). When we combined these genetic perturbations in a *sic1Δnth1Δgph1Δ* strain, Hog1 again adapted in time comparable to WT, supporting the notion that the Gph1-mediated breakdown of glycogen is the main driver of the accelerated *sic1Δ* phenotype (Supplemental Figure 5C). To ensure that the *gph1Δ* rescue is specific to mitigate the effect of *sic1Δ*, and not a broad glycolytic flux perturbation irrespective of genetic background, we tested whether the absence of Gph1-mediated glycogen breakdown affected an otherwise WT Hog1 response. Because glycogen catabolism is halted due to normal cell cycle arrest, *gph1Δ* cells should show a WT adaptation. Indeed, Hog1 adaptation time following osmostress in *gph1Δ* cells is nearly indistinguishable from the WT response (Supplemental Fig 5A, B). Consistent with the rescue in Hog1 dynamics, we observed that the deletion of Gph1 counter-acted the *sic1Δ* effect and reduced the glycerol accumulation at 15 minutes in a manner commensurate to the WT rate (Figure 3D). We attempted to measure cellular glycogen levels and did not observe a significant difference within the first 15 minutes of response between strains (data not shown). This is consistent with the notion that potentially undetectable changes in internal glycogen can be converted to substantial changes in glycerol content, given the massive polysaccharide nature of glycogen and the 3-carbon composition of glycerol.

### Accelerated recovery due to glycogen storage liquidation during osmotic shock prioritizes faster adaptation over robustness to repeated insults

Bypassing cell cycle arrest during osmotic shock results in accelerated recovery due to cell cycle-mediated carbon flux shunted into glycerol production. This suggests that the *sic1Δ* mutant might have an advantage upon one instance of osmotic stress. Conceivably, multiple instances of osmotic shock can amplify the advantage that *sic1Δ* cells have over WT cells. Alternatively, the faster recovery advantage of *sic1Δ* cells might come at the cost of other vulnerabilities that are only revealed dynamically ^35^.

To test the endurance of the *sic1Δ* mutant upon a series of osmotic shocks, we subjected cells to the same 90 minute step input of 1.2 M sorbitol as before, but followed by three 45 minute shocks separated by 5 minutes (Figure 4A). At the end of each step input, we calculated the relative adaptation time using Hog1 nuclear residence as a metric and also assessed the visible physical characteristics of cells. After the first osmotic shock, both strains adapt as previously shown, with the *sic1Δ* mutant recovering faster than its WT counterpart (Figure 4B). Following the first step input, Hog1 adaptation to subsequent osmostress inputs proceeded faster (around 50% faster in this case) in both WT and mutant strains due to accumulation of glycerol from the previous cycle and consistent with previous reports ^35^ (Figure 4C). However, in the subsequent pulses, many *sic1Δ* mutant cells started exhibiting morphological differences from their WT counterparts. These cells displayed a deflated phenotype with visible material accumulated in the nearby vicinity, suggesting the cell lysed and released intracellular debris. This morphology is consistent with a breakdown of cell wall integrity (Supplemental Video), which is a common mechanism of death in serial osmotic shock perturbations ^23^. Assessment of this phenotype revealed a marked increase in cells that have a breakdown in their cell wall integrity for each subsequent pulse in *sic1Δ* cells. By the end of the fourth step input of osmotic shock, 25% of *sic1Δ* cells displayed a morphology consistent with a compromised cell wall, while only 5% of WT cells displayed a similar phenotype (Figure 4D, E). Interestingly, the *sic1Δgph1Δ* genetic background shares the same acute vulnerability as *sic1Δ* to repeated osmotic pulses despite its adaptation that resembles that of the WT. However, the triple mutant *sic1Δnth1Δgph1Δ* is able to recover its morphology after sequential osmoshocks in a manner that is more similar to the WT (Supplemental Figure 6D). This difference in responsiveness suggests unique roles for the breakdown of trehalose and glycogen that warrant further investigation. Nonetheless, it is clear that while the accelerated adaptation ostensibly provides an advantage in the face of a single step input of osmotic shock, the *sic1Δ* mutant is severely ill-suited for repeated insults.

## Discussion

Alterations in cell cycle dynamics have a fundamental presence in many adaptations to stress, but a mechanistic understanding of its role has long been outstanding. Here we attempted to understand some aspects of the role of cell cycle arrest in the context of the well-studied HOG pathway and associated osmotic stress. Our approach was to decouple the HOG stress program from the cell cycle machinery and monitor the nuclear localization dynamics of HOG master effector (Hog1). This experiment revealed that the stereotyped behavior of Hog1 accelerated when the cell cycle could still proceed during osmoshock. We confirmed that the HOG program was still competent under these circumstances by measuring other canonical hallmarks of osmoshock recovery - internal glycerol content and cellular volume. Observing that both proceed faster in the mutant, we utilized quantitative mass spectrometry to implicate internal glycogen liquidation by the mutant as a route by which glycerol synthesis increases and mediates faster adaptation to the stress. In strong agreement with this insight, deletion of the glycogen catabolism enzyme Gph1 in a *sic1Δ* background rescues the Hog1 translocation and glycerol accumulation phenotypes. Intrigued by the observation that the WT is not optimized with respect to the speed of its recovery, we hypothesized that the mutant, which has faster dynamics, might have vulnerabilities that the WT can circumvent. Following this reasoning, we identified a critical failure mode of the mutant by subjecting it to multiple step inputs of osmotic shock. The mutant adapts faster and recovers its basal morphology after the first osmotic shock, but trades its faster initial adaptation for a compromised cell wall in subsequent osmoshocks. Meanwhile, the WT adapts slower after the initial osmotic shock, but maintains nearly 95% consistent physical traits throughout the experiment, thus highlighting the dichotomy between apparent short-term gain versus long-term resilience against a dynamic environment.

We believe that the main contribution of this work is two-fold. First, our investigations provide a higher resolution dissection of the interconnection between three crucial cellular pathways: the cell cycle, the HOG stress response, and carbon metabolism. Contrary to expectation, this connection is not perfectly tuned to maximize the speed of adaptation to stress. In fact, the connection of the HOG pathway to the cell cycle diminishes the ability of the cell to recover rapidly following osmostress, and ablation of this connection allows the cell to recover a substantial 30% faster. The cell cycle seems to be the gatekeeper of a metabolic valve that can augment carbon flow into glycerol from internal resources, but this valve remains shut in WT cells. Opening of this valve in mutant (*sic1Δ*) cells seems to mediate their faster recovery, as evidenced by a *gph1Δ* mutant in which deletion of the enzyme presiding over the internal flux from glycogen to glycerol abolishes the fast osmostress recovery. Therefore, our data provide additional mechanistic details to an intricate interplay of pathways that together set the cellular recovery tempo.

The second contribution of this work is to formulate an instance in which cells seem to navigate a delicate functional balance, sacrificing the brief advantage of faster recovery from an insult for robustness to future environmental changes (Figure 4F). This is perhaps a prompt to revise our view of how to interpret the measured dynamics of stress responses and our assumptions about how cells mobilize their resources to combat stress. It is clear that at least in the example of osmostress, *S. cerevisiae* cells do not maximally mobilize their internal carbon sources to combat the stress, and hence sacrifice substantial speed in their recovery. It is also evident that the cell cycle serves as an arbiter and enforcer of this suboptimal performance. Since the mutant that evades the speed limitation shows tremendous vulnerability to repeated stress, one is compelled to at least hypothesize that this cell cycle control has evolved to alleviate such vulnerability in an environment where repeated or oscillating stress might be more probable.

The work presented here reframes cell cycle arrest in a mechanistic light as being a mediator of a slower adaptation response to hyperosmotic shock. In future investigations, it would be interesting to use a similar approach with conditional or inducible mutants to test for the role of cell cycle arrest in other stresses, potentially discovering similar metabolic flux control mechanisms or roles more tailored to specific stresses. Alternatively, expanding the scope of this question to higher eukaryotes could further illuminate the complex relationship between cell cycle, metabolism and stress response, which has been implicated in several pathologies ^36^. More broadly, it is clear that as we begin to explore how multiple pathways collaborate to allow a cell to navigate its complex environment, we need to revisit statements about functional allocations and re-explore plausible but exceedingly simple assignment of roles and assumptions of unifunctional optimality of any one pathway. We hope that the data presented in this work help to form a basis for such investigations, initiated by our quantitative exploration of the ubiquitous role that cell cycle arrest plays in stress adaptation.

## Materials and Methods

### Strain Construction

The base *S. cerevisiae* strain used in this study is w303. Hog1-mVenus at the endogenous locus was generated by ordering oligos of 40 bp homology 5’ upstream and 40 bp homology downstream of the stop codon, PCR amplifying the mVenus-HIS3 cassette, and transforming as previously described^37^. To knockout genes, 80 bp of homology 5’ to the start codon and 3’ of the stop codon was used to PCR amplify a selection marker cassette and transformed as described above. PCR products using oligos in the 5’ UTR and internal to the selection cassette were used to verify knockouts and insertions.

| Name | Background | Genotype | Description |
| --- | --- | --- | --- |
| yARB001 | w303a | *HOG1-mVenus-HIS3* | "Wild type" strain |
| yARB002 | w303a | *HOG1-mVenus-HIS3, SIC1::TRP1* | *sic1Δ* strain |
| yARB003 | w303a | *HOG1-mVenus-HIS3, SIC1::TRP1, NTH1::NAT* | *sic1Δnth1Δ* strain |
| yARB004 | w303a | *HOG1-mVenus-HIS3, SIC1::TRP1, GPH1::LEU2* | *sic1Δgph1Δ* strain |
| yARB005 | w303a | *HOG1-mVenus-HIS3, SIC1::TRP1, GPH1::LEU2, NTH1::NAT* | *sic1Δnth1Δgph1Δ* strain |
| yARB006 | w303a | *HOG1-mVenus-HIS3, GPH1::LEU2* | *gph1Δ* strain |

### Growth Conditions

Single colonies were picked and inoculated from auxotrophic SD (6.7 g/L Bacto-yeast nitrogen base without amino acids, 2 g/L complete amino acid mix, and 20 g/L dextrose) agar plates. SDC liquid media used throughout the study consisted of 6.7 g/L Bacto-yeast nitrogen base, 2 g/L complete amino acid mix, and 20 g/L dextrose.

### Microscopy

Time-lapse microscopy was collected on a Nikon Ti inverted scope 40x air objective, with Sutter XL lamp illumination and a Hamamatsu Flash 4.0 camera. YFP (515 ex/528 em) channel was collected using a Chroma CFP/YFP filter set with an exposure time of 300 ms. Automated image acquisition was controlled by Nikon NIS Elements proprietary software. The CellAsics Onix2 Microfludic platform was used to control the changing of media. Pressure of the media perfusion was held constant at 10.8 kPa in a microfluidic plate designed to trap haploid yeast (Millipore). To ensure that the yeast cells adapted to conditions within the microfluidic chamber, cells were perfused with normal SDC media for 90 minutes prior to the osmotic shock in all experiments.

### Image Processing

The nucleus of each cell was defined as the mean pixel intensity of the brightest 5% of pixels over the segmented cell in the YFP channel. The remainder of the segmented cell outside the brightest 5% was defined as the cytoplasm. Cell tracking and quantification of nuclear/cytoplasmic enrichment was done using automated yeast cell tracking software implemented in Matlab ^38^. Nuclear enrichment is plotted as the population average nuclear to cytoplasmic ratio divided by the average three time points before the onset of osmotic shock minus one, as previously described ^18^. Cellular volume was calculated from segmentation of an out of focus brightfield image using the Nucleaizer web interface (www.nucleaizer.com). The surface area of each cell in pixels was then converted into an approximate volume as described previously ^18^. This volume was normalized by the three time points before the onset of osmotic shock.

### Mass Spectrometry

Samples were grown overnight to saturation and diluted in SDC media. Cultures were grown to mid-log and 1 mL was transferred to a 0.45 μm PVDF membrane filter paper (Millipore), fixed atop a vacuum manifold. Cells were continuously perfused with either normal SDC media or 1.2 M sorbitol in SDC media for the indicated durations (Figure 2A). At the end of the perfusion period, the filter paper containing cells was immediately transferred to a 2 mL quenching solution of 40:40:20 Methanol:Acetonitrile:H_2_O chilled to −20 °C. After 2 hours incubation at −20 °C, quenching solution plus cells was transferred to a conical tube and dried for approximately 7 hours *in vacuo* and resuspended in 45 μL H_2_O.

In the experiment described in Figure 2C, five minutes prior to time zero 50 mL culture were transferred to a conical tube, spun down at 2000 RPM for 2 minutes and resuspended in 25 mL fully labeled ^13^C glucose SDC. After time zero, at the indicated time points, 1 mL of culture was transferred to the same filter paper vacuum manifold described above, and after media washed through the sample was quenched as described above.

Collected compounds were analyzed using an LC-MS/MS mass spectrometer system consisting of a 1290 Infinity LC (Agilent Technologies) coupled to a 5500 QTRAP triple quadrupole mass spectrometer (AB Sciex) in negative mode and with multiple reaction monitoring (MRM) scan type. Five μL of metabolite extracts were injected on an Agilent PoroShell 120 HILIC-Z column (150 × 2.1 mm, 2.7 μm; Agilent, Santa Clara, CA) using a mobile phase A (water, 10 mM ammonium acetate, 5 μM medronic acid, pH 9) and mobile phase B (90% acetonitrile, 10% water, 10 mM ammonium acetate, 5 μM medronic acid, pH 9) at a constant flow rate of 250 μl/min; Initial conditions: 10% A, 2 min: 10% A, 12 min: 40% A, 15 min: 40% A, 16 min: 10% A, 24 min: 10% A. The MRM settings were adapted from Yuan et al ^39^. The raw data were processed and analyzed using custom software in Matlab (Mathworks), and ^13^C fractional labeling was corrected for natural abundance of ^13^C isotopes as previously described ^40^.

### Intracellular glycerol assay

Each strain was inoculated into SDC media, grown overnight at 30 °C, split and diluted into six 600 μL 0.1 OD600 cultures in a 96 well 2 mL plate. Once the cells were in log phase growth, 600 μL of 2.4 M sorbitol was added to one well at time points of 60 minutes, 30 minutes, 15 minutes, 10 minutes, 5 minutes and 0 minutes. After time zero, 200 μL of each culture was transferred to a separate Corning 3904 96-well assay plate plate for an OD600 reading, and the remaining 1 mL immediately spun down for 2 minutes at 2000 RPM. Cells were washed in 400 μL H_2_O, and spun down again for 2 minutes at 2000 RPM. The culture was then resuspended in 150 μL H_2_O, and left to incubate for 15 minutes at 95 °C. Following incubation at 95 °C, cells were vortexed for 2 minutes and promptly spun down for 10 minutes at 4000 RPM. After the pelleting of cell debris, 100 μL of supernatant was carefully removed and transferred to a separate plate and kept at 4 °C. Colorimetric glycerol assays were acquired using a commercial kit (Sigma) where the provided assay powder was resuspended in 40 mL of distilled H_2_O. For each sample, 5 μL of supernatant was added to 400 μL of glycerol free reagent solution, and left to incubate at room temperature for 15 minutes hidden from light. After 15 minutes, 200 μL of the glycerol free reagent solution-sample mixture was transferred to a separate plate and the OD540 was acquired for each sample on a Tecan Spark 10M plate reader. To account for differences in cell density across samples, the 540 nm readings were normalized by their OD 600 nm reading values.

### Growth Assay

Each strain was inoculated into SDC media overnight, reached saturation and diluted the following morning to an OD600 of 0.1. After dilution, 200 μL of culture was transferred to a Corning 3904 96-well assay plate and grown at 30 °C while shaking. Optical density readings were collected at 600 nm every 20 minutes until saturation on a Tecan Spark 10M plate reader.

### Cell Morphology Quantification

At the end of each 45 minute step input of osmotic shock, each cell was manually assessed for cell cycle re-entry and return to basal morphology. Cells that either did not show continued cell cycle progression, or displayed a visible change in refractive index reflective of a change in morphology were labeled with altered morphology.

## Author Contributions

A.R.B. and H.E-.S. conceived of the study. A.R.B., K.K, and M.D. collected and processed data. A.R.B., K.K., and H.E-.S. interpreted results. A.R.B. and H.E-.S. wrote and edited the manuscript with input from all authors.

## Acknowledgements

The authors thank the members of the El-Samad lab, Joe DeRisi (UCSF) and Sophie Dumont (UCSF) for helpful feedback and discussions. The authors also thank Uwe Sauer (ETH Zürich) for reagents, and DeLaine Larsen and Kari Herrington (UCSF Nikon Imaging Center) for microscope assistance. H.E-.S. is an investigator in the Chan Zuckerberg Biohub and this work was supported by the CZ-Biohub gift. This work was also supported by NIH grant R01GM119033 awarded to H.E-.S. and the National Defense Science & Engineering Graduate (NDSEG) Fellowship awarded to A.R.B.

## Notes

### Competing Interest Statement

The authors have declared no competing interest.

