## Supplemental Figures and Table for "Stress-Induced Transient Cell Cycle Arrest Coordinates Metabolic Resource Allocation to Balance Adaptive Tradeoffs"

**Supplemental Figure Captions:**

**Supplemental Figure 1: The *sic1Δ* mutant recovers faster following osmotic shock induced by 0.6 M NaCl.** A) Time traces of Hog1-mVenus nuclear enrichment for WT (blue) and *sic1Δ* (orange) in response to a 0.6 M NaCl step input. Shaded regions represent the SEM of n=3 biological replicates. B) Quantification of Hog1 adaptation for WT (blue) and *sic1Δ* (orange). Values are normalized to the average WT. Error bars represent the SEM of n=3 biological replicates. *P<0.05; two-sided Student's *t*-test.

**Supplemental Figure 2: External $^{13}C$ incorporation rate in WT is greater than *sic1Δ* by nearly three-fold in the presence and absence of osmotic shock.** A) Cartoon schematic of experiment to infer extracellular glucose incorporation rates. At time zero a 1 mL sample of cells was transferred to filter paper above a vacuum manifold and continuously perfused with fully-labeled $^{13}C$ glucose media with and without 1.2 M sorbitol for durations of 10 s, 20 s, 30 s, 45 s, and 60 s and transferred to quenching solution. B) A schematic of central glycolysis and the glycerol branch of central metabolism with the targeted metabolites featured. WT (blue) and *sic1Δ* (orange); solid lines represent 1.2 M sorbitol osmotic shock, and dashed lines represent normal defined media. Error bars represent the standard deviation of n=2 technical replicates. C) Using the average total ion count ($^{12}C$ + $^{13}C$) and the exponential decay constant of each $^{12}C$ enrichment for each metabolite, bar plots show the average flux approximation per condition. The plotted value represents the average of the n=2 technical replicates (Supplemental Table 1).

**Supplemental Figure 3: Average growth rate bears a weak correlation to average Hog1 adaptation time.** A) OD600 readings of WT (blue), *sic1Δ* (orange), *sic1Δnth1Δ* (grey), *sic1Δgph1Δ* (purple) ,*sic1Δnth1Δgph1Δ* (gold) and *gph1Δ* (green) strains with readings taken every 20 minutes over 24 hours. Shaded regions represent the SEM of at least n=2 biological replicates. B) The average growth rate of each strain measured in Panel A plotted against its average adaptation time later shown in Supplemental Figure 5B. Error bars along each axis represent the standard deviation of at least n=2 biological replicates. R, Pearson's correlation coefficient. P-value; Student's t-test.

**Supplemental Figure 4: Stress-induced mobilization of an internal carbon macromolecule is shunted into central glycolysis in the *sic1Δ* mutant.** A) Cartoon schematic of experiment to test internal carbon enrichment of targeted metabolites. Five minutes prior to time zero, cells were resuspended in fully-labeled $^{13}$C glucose. At time zero the culture of cells were diluted 1:1 in fully-labeled in $^{13}$C glucose with either 2.4 M sorbitol or normal defined media. At the indicated time points, 1 mL of culture was placed on filter paper above a vacuum manifold for the media to wash through, and transferred to quenching solution. B) A schematic of central glycolysis and the glycerol branch of central metabolism with the targeted metabolites featured. WT (blue) and *sic1Δ* (orange); solid lines represent 1.2 M sorbitol osmotic shock, and dashed lines represent normal defined media. Error bars represent the standard deviation of n=2 technical replicates.

**Supplemental Figure 5: Knockout of glycogen catabolism enzyme, Gph1, is sufficient and necessary to rescue accelerated phenotype conferred with removal of Sic1.** A) Traces of Hog1 nuclear enrichment over time following 1.2 M sorbitol osmotic shock in the WT (blue), *sic1Δ* (orange), *sic1Δnth1Δ* (grey), *sic1Δgph1Δ* (purple) ,*sic1Δnth1Δgph1Δ* (gold) and *gph1Δ* (green) cells. Shaded regions represent the SEM of n=3 biological replicates. B) Quantification of adaptation time of Hog1 nuclear enrichment computed as in Figure 1B. Values are normalized to the average WT. Shaded regions represent the SEM of n=3 biological replicates. *$P<0.05$; two-sided Student's *t*-test. C) Quantification of internal glycerol as a function of time for WT (blue), *sic1Δ* (orange), *sic1Δnth1Δ* (grey), *sic1Δgph1Δ* (purple) ,*sic1Δnth1Δgph1Δ* (gold) to a step input of 1.2 M sorbitol osmotic shock. Measurements are taken using a colorimetric assay. Error bars represent the standard deviation for n=3 biological replicates. D) The change in glycerol, calculated as the difference between two time points for data in Panel C, is plotted as a function of time. Error bars represent the standard deviation for n=3 biological replicates.

**Supplemental Figure 6: Multiple step inputs of osmotic shock reveal different susceptibilities between rescue strains.** A) Top: experiment schematic representing a series of 1.2 M sorbitol osmotic shock step inputs. The first input lasts for 90 minutes and subsequent inputs last 45 minutes, and are separated by 5 minutes. B) Time traces of Hog1 nuclear enrichment of WT (blue), *sic1Δ* (orange), *sic1Δgph1Δ* (purple), *sic1Δnth1Δgph1Δ* (gold). Shaded regions represent the SEM of n=3 biological replicates. C) Quantification of adaptation time of Hog1 nuclear enrichment for WT and *sic1Δ* strains for data presented in Panel B. Values are normalized to the average first response for the WT strain. Error bars represent the SEM of n=3 biological replicates. D) The percent of cells with altered morphological phenotypes at the end of each step input of 1.2 M sorbitol. Error bars represent the SEM of n=3 biological replicates. **$P<0.005$; two-sided Student's *t*-test.

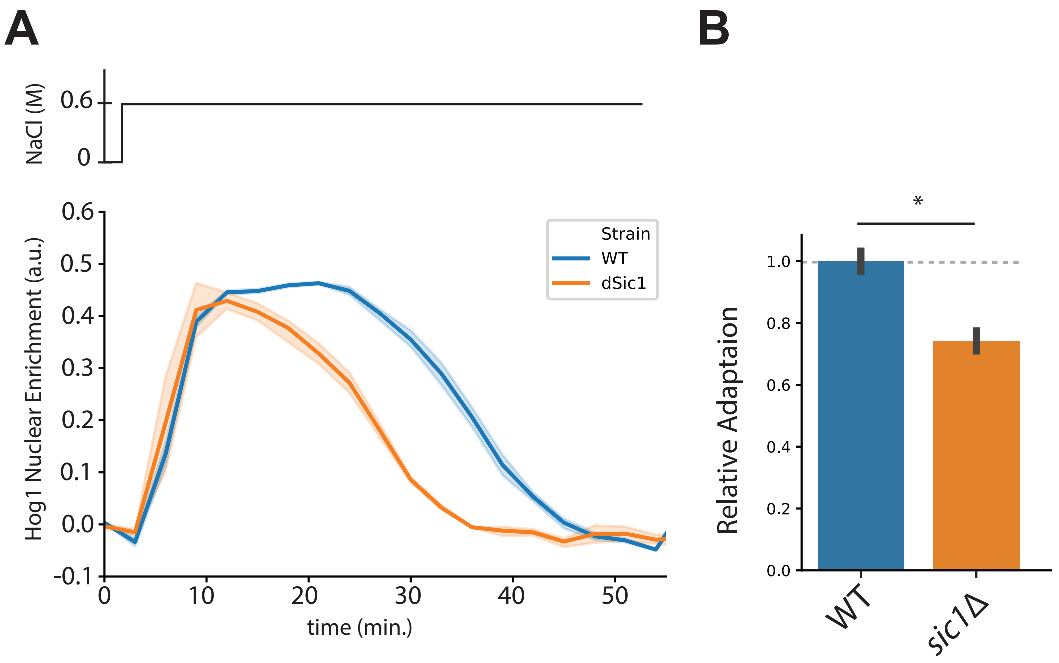


Supplemental Figure 1

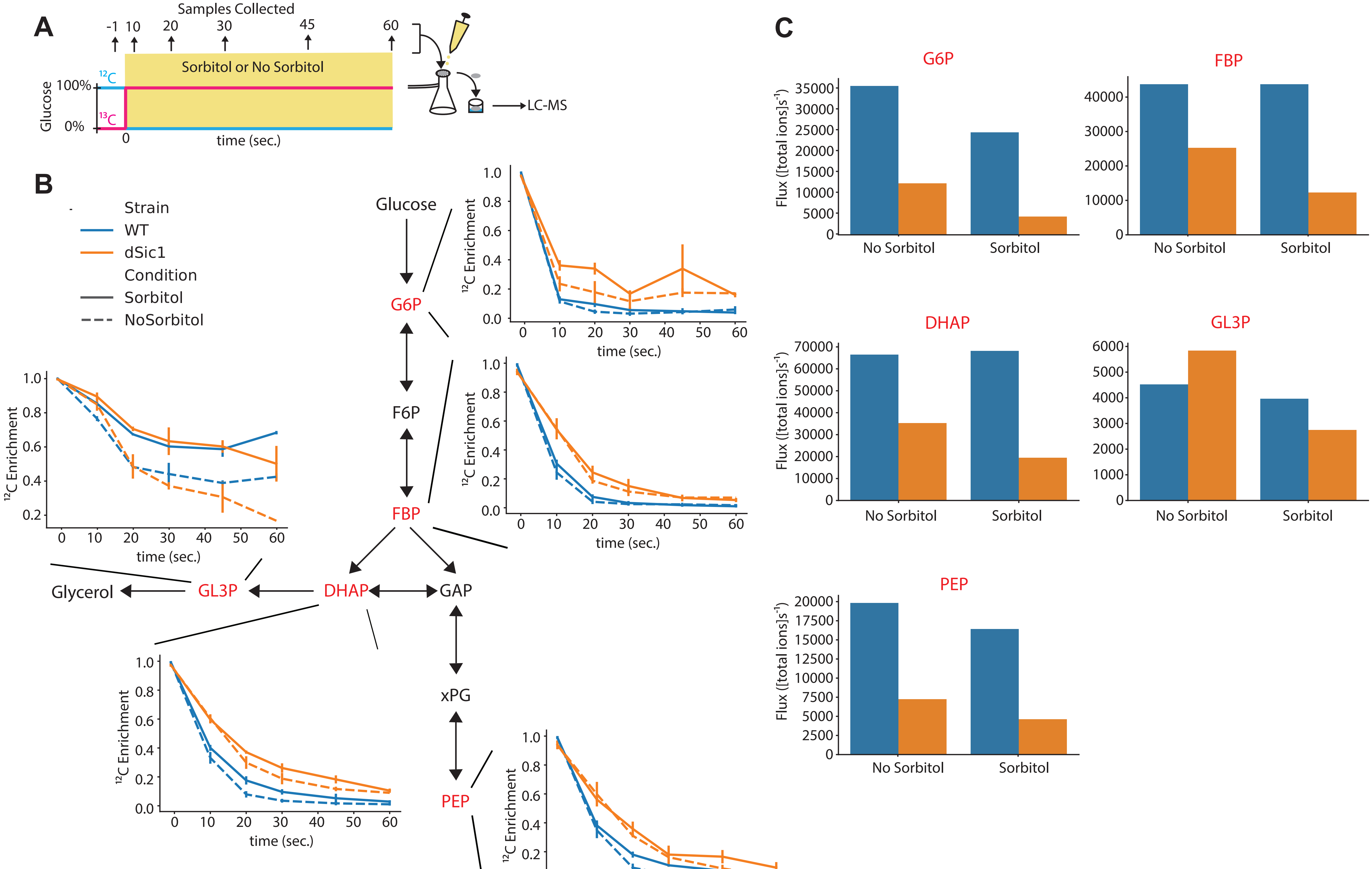


Supplemental Figure 2

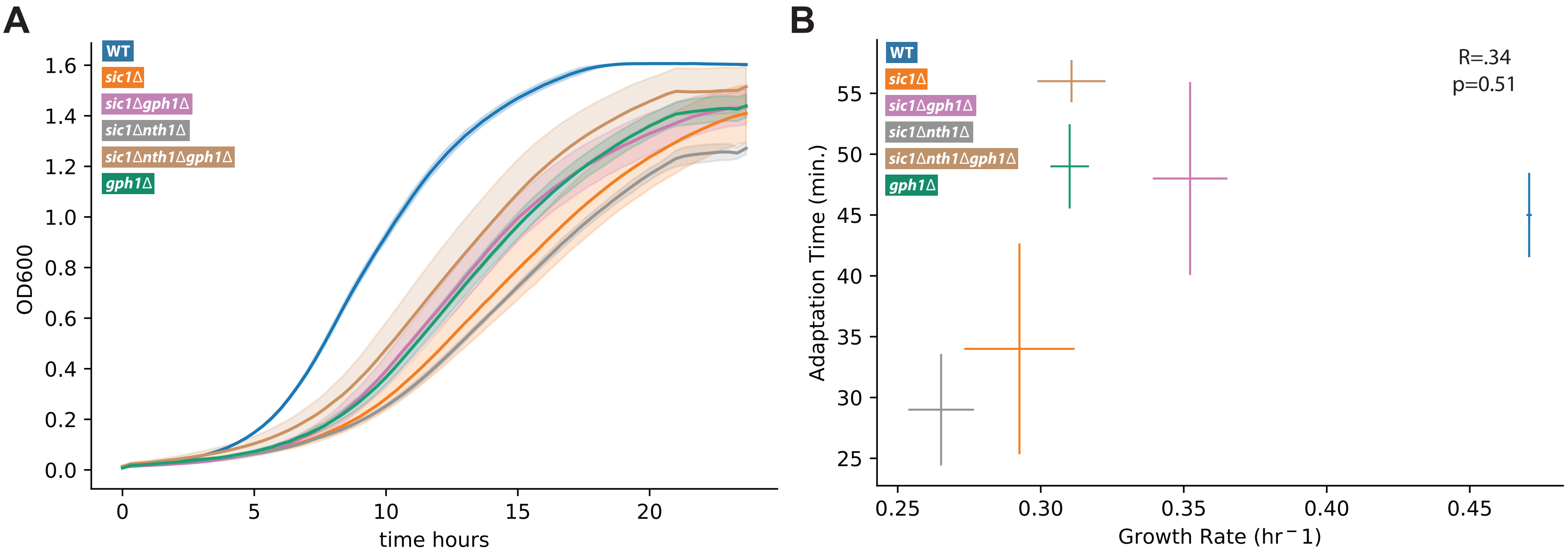


Supplemental Figure 3

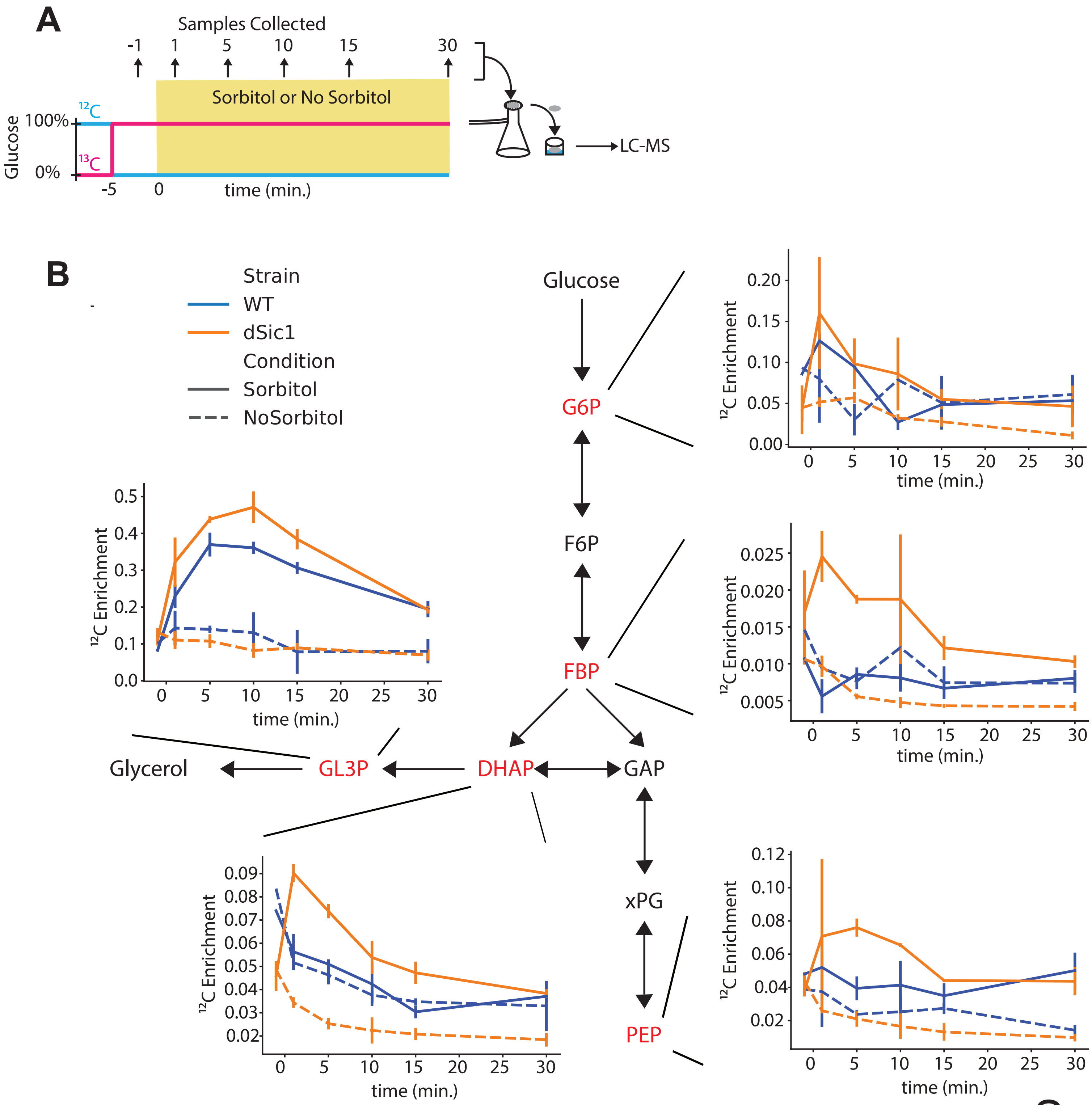


Supplemental Figure 4

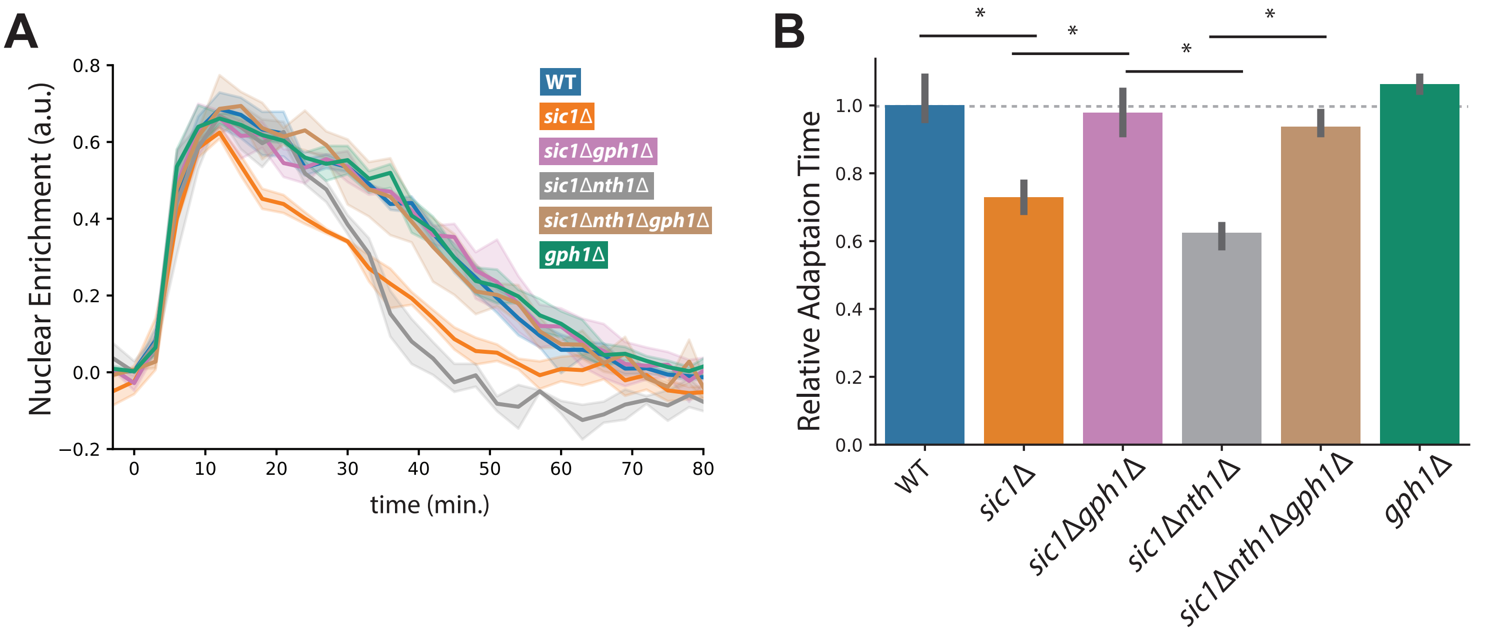


Supplemental Figure 5

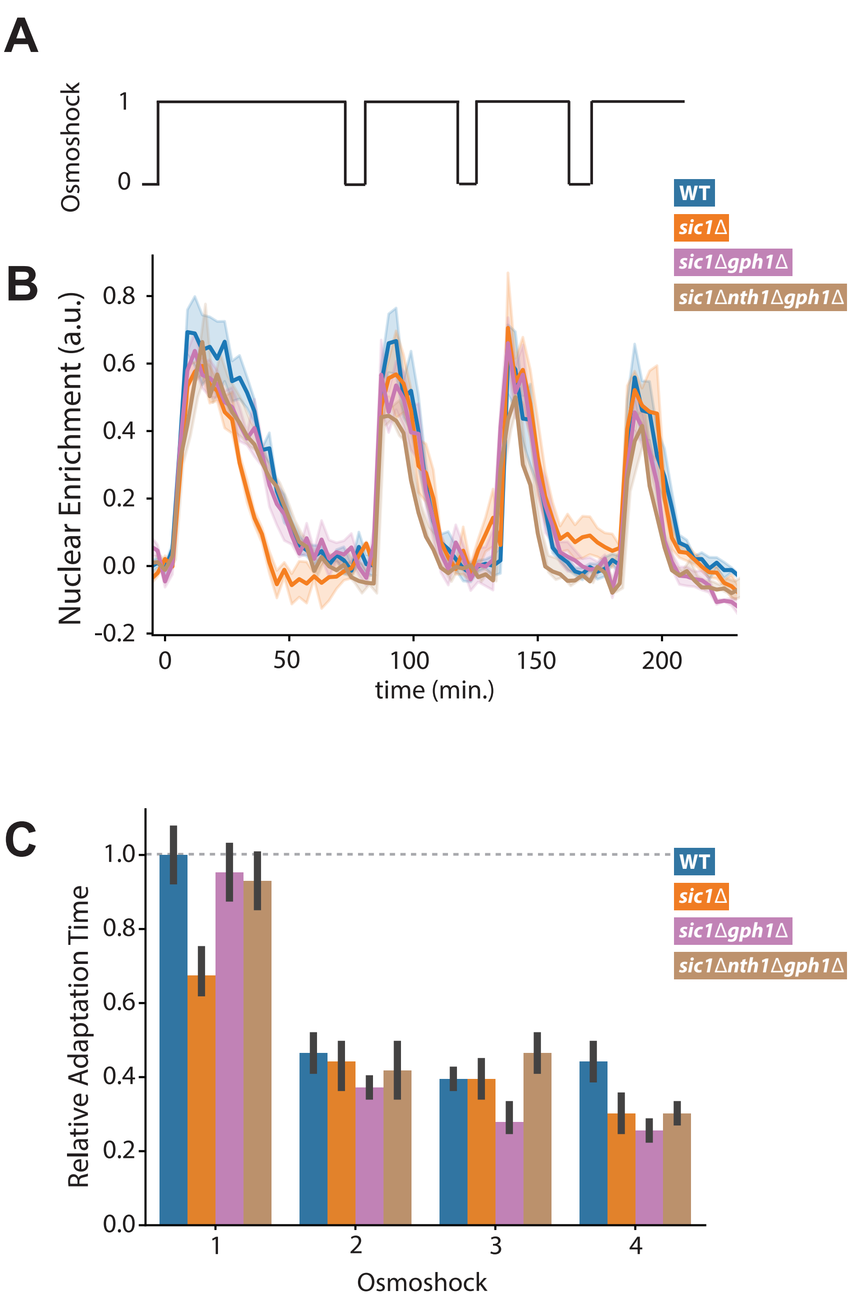


Supplemental Figure 6

| Metabolite | Condition | Strain | Decay Rate | Incorporation Rate | Total Ions | Flux Approximation |
|---|---|---|---|---|---|---|
| FBP | NoSorbitol | WT | -0.15 | 0.15 | 2914547.5 | 437182.125 |
| FBP | Sorbitol | WT | -0.12 | 0.12 | 3640760.333 | 436891.24 |
| G6P | NoSorbitol | WT | -0.16 | 0.16 | 222888.5 | 35662.16 |
| G6P | Sorbitol | WT | -0.117 | 0.117 | 209617.75 | 24525.27675 |
| DHAP | NoSorbitol | WT | -0.103 | 0.103 | 644957.75 | 66430.64825 |
| DHAP | Sorbitol | WT | -0.075 | 0.075 | 909824.5 | 68236.8375 |
| PEP | NoSorbitol | WT | -0.105 | 0.105 | 188753.25 | 19819.09125 |
| PEP | Sorbitol | WT | -0.0793 | 0.0793 | 206968.25 | 16412.58223 |
| Glycerol-3-P | NoSorbitol | WT | -0.0345 | 0.0345 | 130922.25 | 4516.817625 |
| Glycerol-3-P | Sorbitol | WT | -0.0187 | 0.0187 | 212167 | 3967.5229 |
| FBP | NoSorbitol | dSic1 | -0.076 | 0.076 | 3321460.75 | 252431.017 |
| FBP | Sorbitol | dSic1 | -0.064 | 0.064 | 1915263.75 | 122576.88 |
| G6P | NoSorbitol | dSic1 | -0.084 | 0.084 | 146522.75 | 12307.911 |
| G6P | Sorbitol | dSic1 | -0.04 | 0.04 | 109059 | 4362.36 |
| DHAP | NoSorbitol | dSic1 | -0.052 | 0.052 | 678069 | 35259.588 |
| DHAP | Sorbitol | dSic1 | -0.0375 | 0.0375 | 518724.25 | 19452.15938 |
| PEP | NoSorbitol | dSic1 | -0.055 | 0.055 | 131545.75 | 7235.01625 |
| PEP | Sorbitol | dSic1 | -0.0515 | 0.0515 | 89478.5 | 4608.14275 |
| Glycerol-3-P | NoSorbitol | dSic1 | -0.0339 | 0.0339 | 172045.75 | 5832.350925 |
| Glycerol-3-P | Sorbitol | dSic1 | -0.0173 | 0.0173 | 158569.5 | 2743.25235 |
| 6PG | NoSorbitol | dSic1 | -0.015 | 0.015 | 21516.5 | 322.7475 |
| 6PG | Sorbitol | dSic1 | -0.0318 | 0.0318 | 20994 | 667.6092 |
| 6PG | Sorbitol | WT | -0.0575 | 0.0575 | 36595 | 2104.2125 |
| 6PG | NoSorbitol | WT | -0.054 | 0.054 | 31091 | 1678.914 |
